# Learning from local to global - an efficient distributed algorithm for modeling time-to-event data

**DOI:** 10.1101/2020.03.04.977298

**Authors:** Rui Duan, Chongliang Luo, Martijn J. Schuemie, Jiayi Tong, Jason C. Liang, Howard H. Chang, Mary Regina Boland, Jiang Bian, Hua Xu, John H. Holmes, Christopher B. Forrest, Sally C. Morton, Jesse A. Berlin, Jason H. Moore, Kevin B. Mahoney, Yong Chen

**Author notes:** Co-first authors (Rui Duan, Chongliang Luo, Martijn J. Schuemie). Correspondence: Yong Chen, PhD, University of Pennsylvania School of Medicine, 423 Guardian Drive, Philadelphia, PA 19104.

## Abstract

**Objectives:** We developed and evaluated a privacy-preserving **O**ne-shot **D**istributed **A**lgorithm to fit a multi-center **C**ox proportional hazard model (**ODAC**) without sharing patient-level information across sites.

**Methods:** Using patient-level data from a single site combined with only aggregated information from other sites, we constructed a surrogate likelihood function, approximating the Cox partial likelihood function obtained using patient-level data from all sites. By maximizing the surrogate likelihood function, each site obtained a local estimate of the model parameter, and the ODAC estimator was constructed as a weighted average of all the local estimates. We evaluated the performance of ODAC with (1) a simulation study, and (2) a real-world use case study using four datasets from the Observational Health Data Sciences and Informatics (OHDSI) network.

**Results:** Our simulation study showed that ODAC provided estimates nearly the same as the estimator obtained by analyzing, in a single dataset, the combined patient-level data from all sites (i.e., the pooled estimator). The relative bias was less than 0.1% across all scenarios. The accuracy of ODAC remained high across different sample sizes and event rates. On the other hand, the metaanalysis estimator, which was obtained by the inverse variance weighted average of the sitespecific estimates, had substantial bias when the event rate is less than 5%, with the relative bias reaching 20% when the event rate is 1%. In the OHDSI network application, the ODAC estimates have a relative bias less than 5% for 15 out of 16 log hazard ratios; while the meta-analysis estimates had substantially higher bias than ODAC.

**Conclusions:** ODAC is a privacy-preserving and non-iterative method for implementing time-to-event analyses across multiple sites. It provides estimates on par with the pooled estimator and substantially outperforms the meta-analysis estimator when the event is uncommon, making it extremely suitable for studying rare events and diseases in a distributed manner.

## INTRODUCTION

Real-world data (RWD) such as the Electronic Health Records (EHRs) and health plan claims are playing an increasing role in generating real-world evidence (RWE) to support health care decision-making [1]. Patient data (e.g., diagnoses, medications, labs, and clinical notes) are routinely collected and entered into the EHRs during clinical care delivery. When compared to cross-sectional observational data, the detailed longitudinal information contained in the EHR enables time-to-event modeling, also known as survival analysis. Incorporating time into statistical models that predict occurrence of outcomes enables a better understanding of the predictors of when an event occurs rather than merely whether it occurred [2–4]. The Cox proportional hazards model (hereafter referred as the Cox model) is one of the most commonly used time-to-event models, and has been widely applied in EHR-based studies for treatment evaluation, risk factor identification and individual risk prediction [5].

The last few years have witnessed an increasing number of clinical research networks (CRNs), curating and using immense collections of health system EHRs and health plan claims data. Two prominent examples are (1) the Observational Health Data Sciences and Informatics (OHDSI) network—an international network of observational health databases that cover more than half a billion patient records in a common data model (CDM), and (2) the national Patient-Centered Clinical Research Network (PCORnet)—a network covering more than 100 million patients in the United States [6, 7]. These large data consortia strive to provide platforms and tools to integrate heterogenous RWD from a diverse range of healthcare organizations. Multicenter analyses using RWD from these CRNs have been increasing rapidly in large part because they can improve the generalizability of the study results by increasing the sample size.

Despite the benefits of multicenter analyses, direct sharing of patient-level data across institutions may be challenging because of concerns related to patient and institutional privacy. Studies that include institutional data from multiple countries face legal and regulatory barriers to sharing patient-level data. The OHDSI network, which is an international consortium, uses a federated model in which the patient-level data are stored at local institutions and only aggregated information are shared. Thus, multicenter analyses have to be based on the summary statistics obtained from each site [8–11]. The results are often combined through meta-analysis, specifically, some form of weighted average where the weight for each estimate, e.g., could be the inverse of the site-specific variance (for a fixed-effect model). However, when the outcomes or exposures are rare, or some of the sites may have small sample size, the accuracy (in terms of bias and precision) of the meta-analysis may be poor, as will be shown later in both the simulation studies and with real world examples.

Distributed computing has been considered in many applications [12–17], where the model estimation is decomposed into smaller computational tasks that are computed locally at each site and then returned to the site instituting the study. For example, a distributed algorithm for conducting logistic regression, GLORE (Grid Binary Logistic Regression) [13], and the WebDISCO (a web service for distributed Cox model learning) for fitting the Cox model [14] were developed and successfully implemented in the pSCANNER (patient-centered Scalable National Network for Effectiveness Research) consortium [18]. Through multiple rounds of iterative communication of information across sites, these algorithms can be lossless (i.e., the results are equivalent to the results from fitting the model on the dataset created by pooling the individual-level data across all sites). However, due to the need to iteratively transfer data across sites, it is time-consuming and labor-intensive in practice. Thus, development of non-iterative distributed algorithms, which require neither data sharing nor iterative communication back and forth between sites, is an active research. For example, Chen et al. developed a lossless non-iterative distributed algorithm for linear regression [12]. Duan et al. proposed a non-iterative distributed algorithm for logistic regression through the construction of a surrogate likelihood [17].

In this paper, we propose the <u>O</u>ne-shot <u>D</u>istributed <u>A</u>lgorithm for <u>C</u>ox model (termed as ODAC). A unique challenge in developing distributed algorithms for the Cox model, compared to our earlier work on logistic regression [17], is that the log partial likelihood function is not equivalent to the summation of local log partial likelihood functions from each site. Each component of the log partial likelihood function involves a non-linear function of data from all patients who are at risk at a certain time point. By designing a novel initialization step, our algorithm extends the surrogate likelihood approach [16] to deal with the more complicated Cox partial likelihood function, while maintaining the property of only requiring communication of aggregated information between sites. We show that our proposed algorithm is non-iterative while achieving high accuracy (small bias) in both a simulation study and a real-world use case that studies the risk factors of acute myocardial infarction (AMI) and stroke using claims data from four different databases in the OHDSI network.

## METHODS

### Cox model

We first introduce the notation and basic assumptions of the Cox model. Let *X* be a vector denoting *p* risk factors and let *T* be the time-to-event for the outcome of interest. The Cox model assumes the hazard at time *t* follows

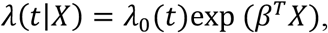

where *λ*_0_(*t*) is the baseline hazard function, and *β* is the vector of regression coefficients which are interpreted as the log-hazard ratios. The observed data are {*T_i_, δ_i_, x_i_*} for the *i*-th subject, with *δ_1_* = 0 indicating censoring and *δ_1_* = 1 indicating an event, *i* = 1, …, *N*. For a given time *t*, we denote *R*(*t*) = {*i*; *T_i_* ≥ *t*}, which is the risk set at time *t*. The log partial likelihood function is constructed as

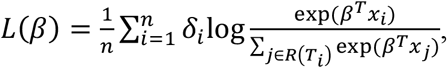

which does not require estimation of the baseline hazard *λ*_0_. The parameter *β* is estimated by the value that maximizes the log partial likelihood function [5].

### Parameter estimation in a distributed network via ODAC

Now suppose we have data stored in K different clinical sites and denote *n_j_* to be the sample size of the data at the *j-th* site. Denote the total sample size to be 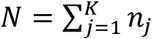. For the *i*-th patient at the *j*-th site, we observe {*T_ij_, δ_ij_, x_ij_*}. The risk set in site *j* at time *t* is defined as *R_j_*(*t*) = {*i; T_ij_* ≥ *t*}, which contains all the subjects in site *j* that have not experienced an event or been censored at time *t*. The combined risk set over all the K sites is *R*(*t*) = {(*i,j*); *T_ij_* ≥ *t*}. With a slight abuse of notation, we further denote *R_ij_* to be the risk set at time *T_ij_*, i.e., *R_ij_* = *R*(*T_ij_*). Denote the set of event times at the *j*-th site to be 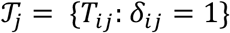, and the total set of unique event time to be 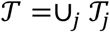. Assume there are in total *d* unique event time points 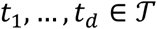. Ideally, if all the data could be pooled together, the combined partial likelihood function could be expressed as

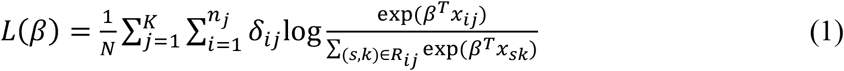

and a pooled estimator 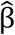 could be obtained by maximizing the combined partial likelihood function. However, in reality, transferring patient-level data across sites is almost always impossible due to patient privacy concerns; thus, each site can only access their own data.

Inspired by the surrogate likelihood approach developed in [16] and [19], we aim to construct an proxy of the combined partial likelihood function, which we call a surrogate likelihood function. The ideal is to construct a function that is close to the combined partial likelihood function defined in equation (1) near a neighborhood of the true parameter value. One naive choice of the surrogate likelihood function (for exampe, at the *j*-th site) is the local partial likelihood function *L_j_*(*β*) constructed as

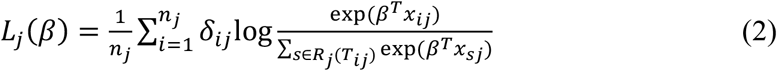

which only utilizes the patient-level data at the *j*-th site. However, this approximation does not utilize any information from other sites. Our goal is to further improve the accuracy of the approximation by borrowing aggregated information from other sites.

Define 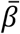 to be an initial value that is in a neighborhood of the true value of the parameter *β*. We propose the surrogate likelihood function obtained at site *j* to be

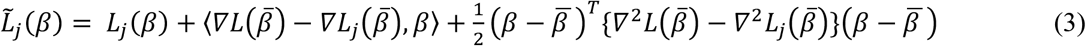

for *j* = 1, … *K*, where *L_j_*(*β*) is the local likelihood function defined in equation (2), *∇* and *∇*^2^ denote the first and second order gradients of a function (explicit forms of 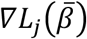, 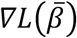, 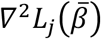 and 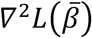 can be found in the Supplementary Material). Intuitively, the surrogate likelihood function in equation (3) utilizes the local likelihood function *L_j_*(*β*) as a baseline function, and it also adds a first-order term and a second-order term, 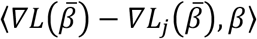 and 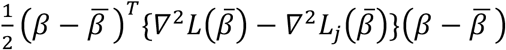, to alter the shape of the local likelihood function arround 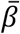 towards the combined likelihood function.

In the construction of the surrogate likelihood function 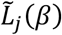, once 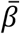 is obtained, the terms 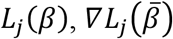 and 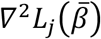 can be calculated at the *j*-th site without needing to transfer information. And the constructing componenets 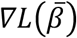 and 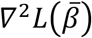 can be calculated distributively from all sites with the help of some preliminary summary-level statistics 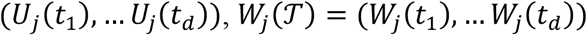 and 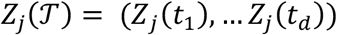, where for each time point *t* in 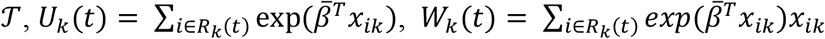, and 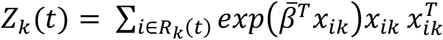. The detailed steps of distributively calculating 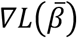 and 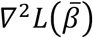 can be found in the Supplementary Material. Importantly, these gradients are all aggregated such that patient privacy is protected.

Regarding the initial value 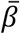, we recommend using the inverse variance weighted average of the estimates obtained by fitting a Cox model at each site, i.e.,

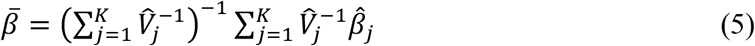

where 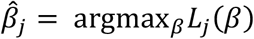 is the estimator from of Cox model fitted on data at the *j*-th site, and *V_j_* is the estimated variance of 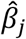. After constructing the surrogate likelihood function at each site, we can obtain a surrogate likelihood estimator at each site by

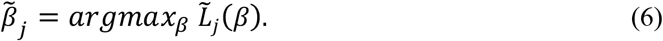

The variance of 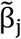 is defined to be 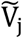, and it can be calculated using Equation S.5 in the Supplementary Material. Finally, we combine all the local surrogate estimator 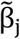 using inverse variance weighted averaging in the same format as in equation (5). We summarize the ODAC algorithm below and also provide a schematic illutration in Figure 1.

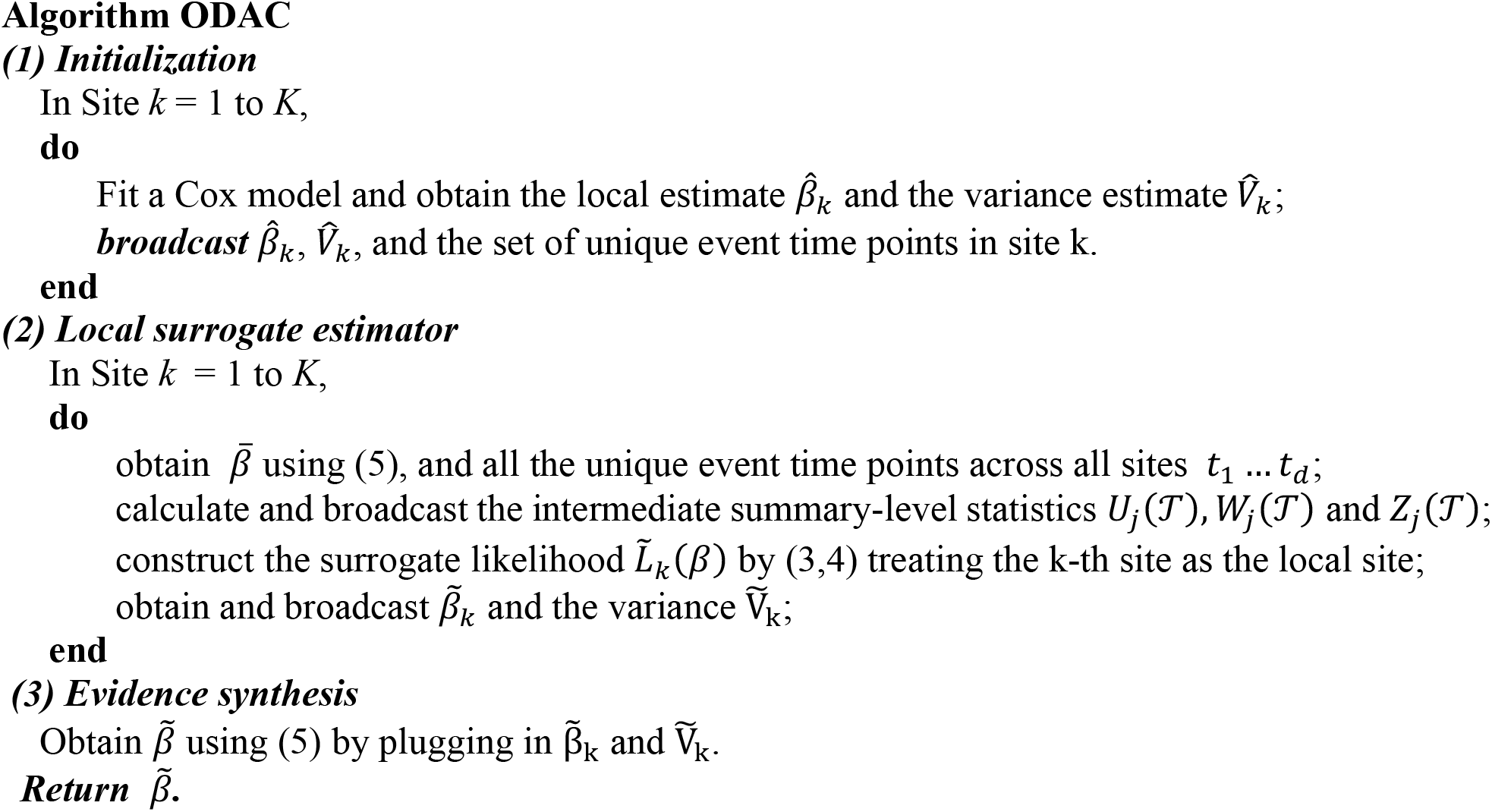

**Figure 1:**
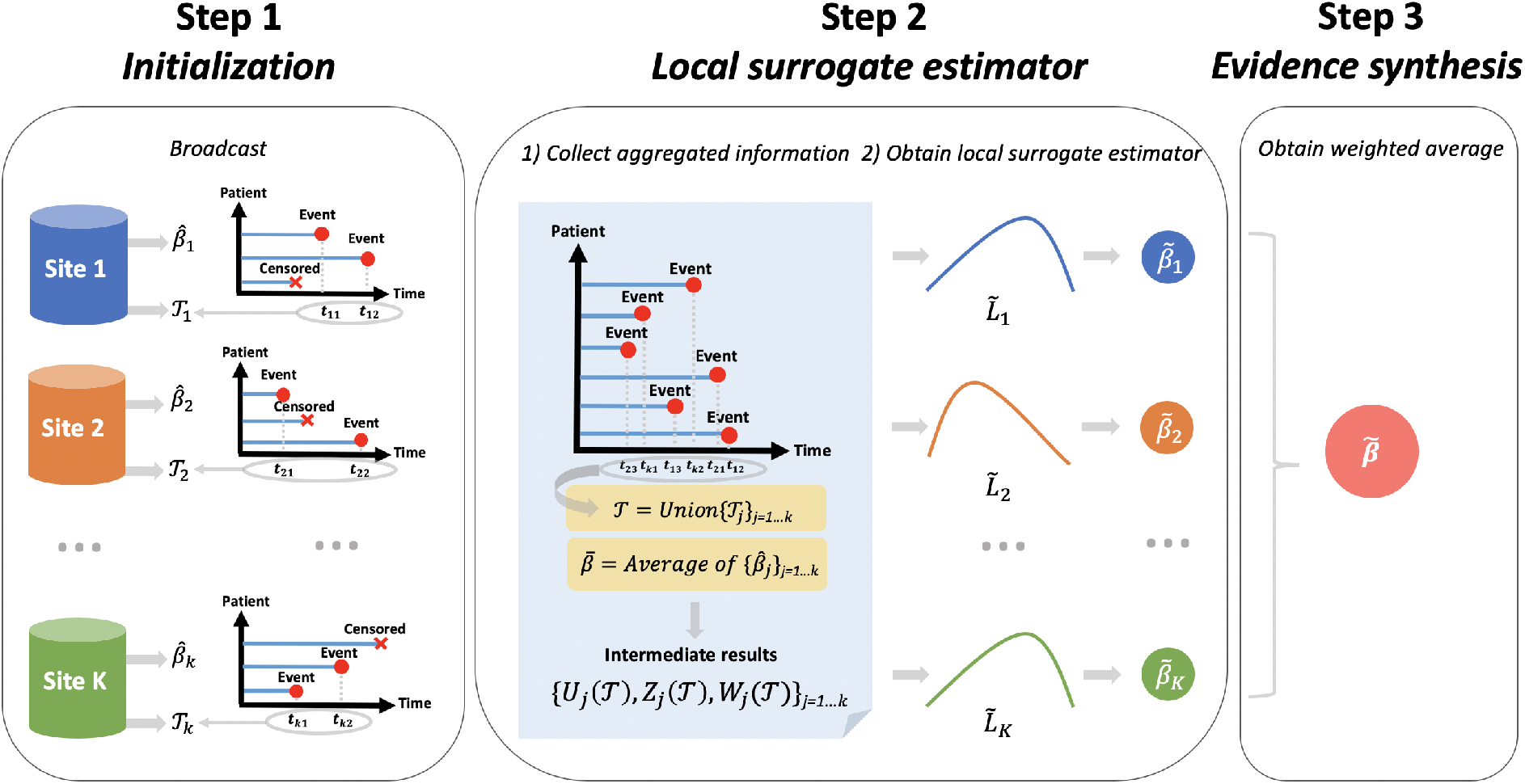
Schematic illustration of the ODAC algorithm. The first step is initialization where each site reports the local estimate of the log hazard ratio 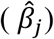, and the set of event times 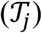. In the second step, each site calculates the average of all local estimates, and obtain the union of all event times. Using this information, each site shares the intermediate results 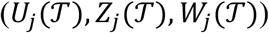. Next, each site combines the intermediate results and the local patientlevel information to construct a surrogate likelihood function defined in equation (3), and obtains the local surrogate estimates by optimizing the surrogate likelihood function. The last step is the evidence synthesis, where all the local surrogate estimates are combined through a weighted average.

### A step-by-step illustration of ODAC using a simulated network with two sites

To better explain each step in the ODAC algorithm, we simulate two datasets that mimic a distributed network with two clinical sites. Site A has 100 patients and Site B has 50 patients, and the goal is to fit a Cox model to study the association between the survival time and treatment (new drug vs placebo) adjusting for age. We generated age from a uniform distribution with a range from 20 to 60, and the treatment status was generated from a Bernoulli distribution with probability 0.5 of being in each arm. Before fitting the model, we standardized age by subtracting the mean age and dividing by the standard error. The survival time of each patient was then generated from a Weibull proportional hazard model, where the baseline hazard follows a Weibull-distribution with scale 200 and shape 20 and the regression parameters for the treatment option and age are set to −1 and 1, respectively. We generated non-informative censoring from a Weibull-distribution to keep the event rate at approximately 5%. **Figure 2** provides a detailed step-by-step illustration of the implementation of the ODAC algorithm to this simulated dataset.

**Figure 2:**
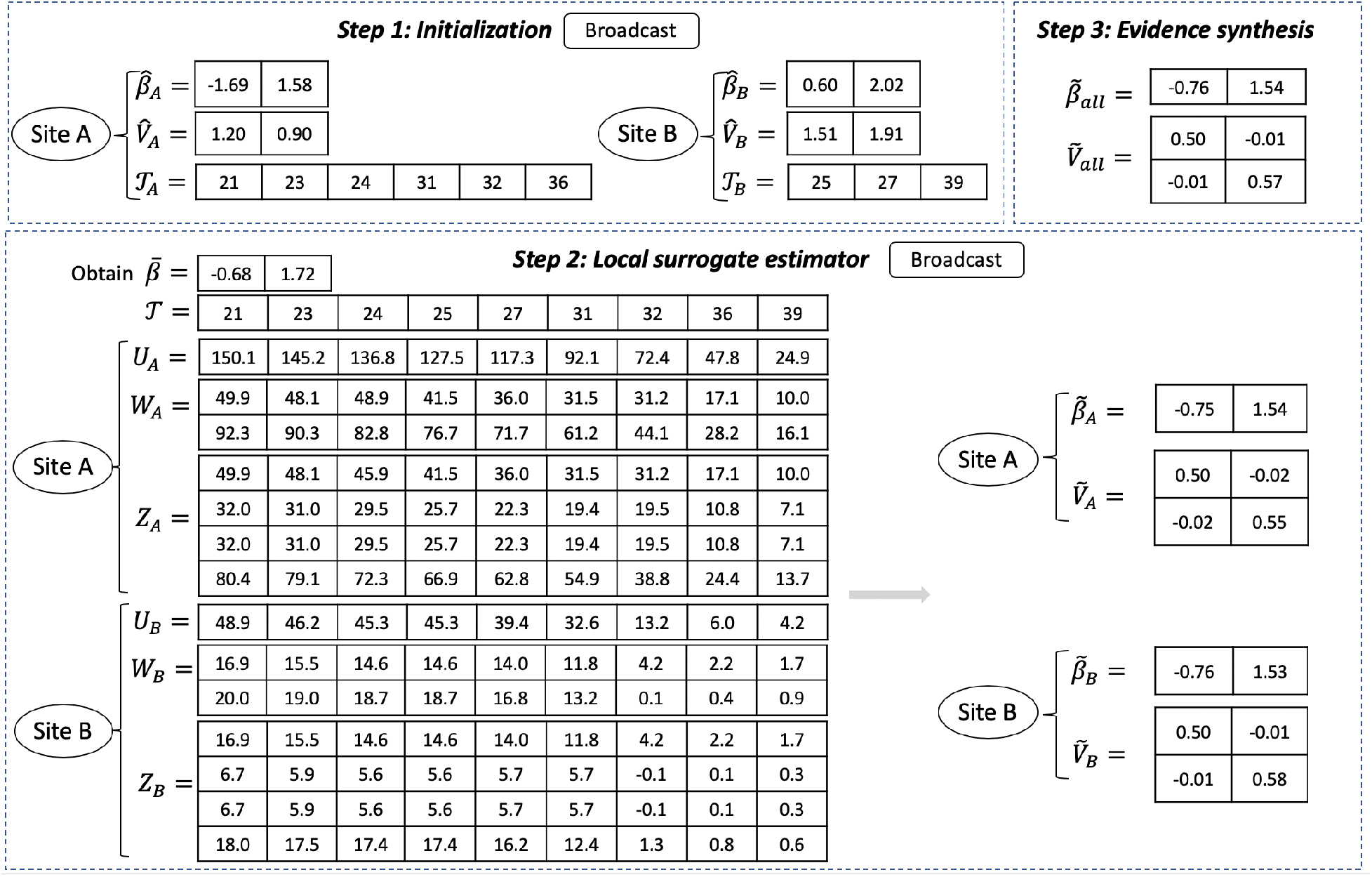
Step-by-Step illustration of the ODAC algorithm using a simulation dataset. In the first step, the two sites share the estimated log hazard ratios and their variances 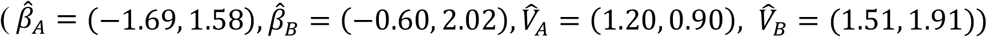 as well as their set of unique event times 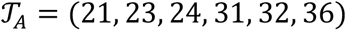, and 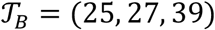. In the second step, each site calculates the inverse variance weighted initial estimator 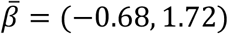, and obtains the combined set of event times 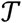 which contains nine unique time points. Then each site shares 63numbers in this step. Using these numbers, Site A and B obtain and share the surrogate likelihood estimates 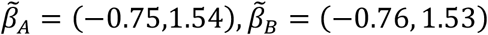, as well as the covariance matrix 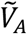 and 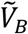. In the evidence synthesis step, the final estimator 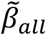 is obtained by the weighted average of 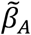 and 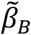 where 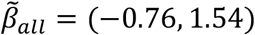.

In the first step, the two sites fit the Cox model locally and share the initial estimates of the log hazard ratios and their variances 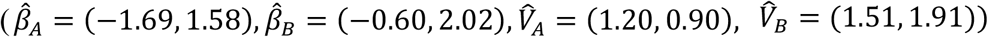. In addition, each site reports their set of unique event times. In Site A, there are six time points at which an event occurs, i.e. 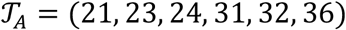, and in Site B we have 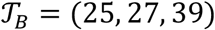. Therefore, in the first step, Site A shares 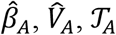 and Site B shares 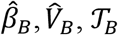.

In the second step, each site calculates the inverse variance weighted initial estimator 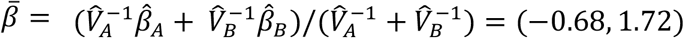, and obtains the combined set of event times 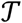 which contains nine unique time points. Then for each time 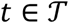, Site A calculates a scalar *U_A_*(*t*), a *p*-dimensional vector *W_A_*(*t*), and a matrix containing *p*^2^ numbers *Z_A_*(*t*) (for simple illustration, we vectorize *Z_A_*(*t*) in Figure 2). Therefore, in this step, in total each site shares 42 numbers that do not contains patient-level information. Using these numbers, Sites A and B are able to construct the surrogate likelihood function within each site, and then obtain and share the surrogate likelihood estimates 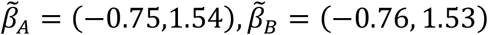, as well as their corresponding covariance matrices 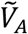 and 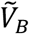.

In the evidence synthesis step, the final estimator 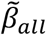 is obtained as the weighted average of 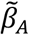 and 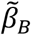 with weights equal to the inverse of 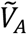 and 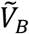. In this example, we obtain 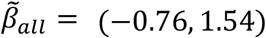, and the variance of 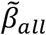 is estimated by 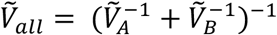, which is closer to the true parameter value than the local estimates.

### A real-world use case using OHDSI Data

We used ODAC to study the risk factors of acute myocardial infarction (AMI) and stroke in a population with pharmacologically-treated major depressive disorder using data from four different US insurance claims databases that have been converted to the the Observational Medical Outcomes Partnership (OMOP) CDM [20] in the OHDSI network. Both AMI and stroke were defined as the occurrence of the respective diagnosis codes in an inpatient or emergency room setting. We only counted the first occurrence of each condition per patient in order to preserve independence.

We fit Cox regression models for the two outcomes and corresponding risk factors. For AMI we included known observed risk factors: age, gender, alcohol dependence, hyperlipidemia, hypertensive disorder, depression, obesity, and type II diabetes [21] [22]. Similarly, for stroke the following known risk factors were included: congestive heart failure, coronary arteriosclerosis, hyperlipidemia, ischemic heart disease, renal failure, hypertensive disorder, transient cerebral ischemia, and type II diabetes [23]. We compared our ODAC estimator with the pooled estimator and the estimator from meta-analysis.

### A numerical experiment to demonstrate the benefit of ODAC in studying rare events

To demonstrate the benefit of ODAC comparing to meta-analysis in study rare events, we designed the following numerical experiment. A pooled dataset of N=10,000 subjects were generated based on Weibull proportional hazard model, where the baseline hazard follows a Weibull-distribution with scale 200 and shape 20. We generated two covariates from i.i.d. uniform distributions and the true log hazard ratios were set to be *β*= (−1, 2). We set the event rate (number of cases over number of subjects) as 20%, 5%, 2% and 1% by appropriately modifying the distribution of censoring times. The pooled data were distributed over K=10 clinical sites, with five large and five small sites. We set the relative sizes of the large to small sites to be 1000/1000, 1250/750, 1500/500. Under each scenario, we compared the ODAC estimator to the meta-analysis estimator over 1000 replications. Since the pooled estimator can be considered a gold standard, the bias to the pooled estimator is used as the metric to evaluate the performance of each method. For simplicity of illustration, we only present the results for estimation of coefficient *β*_2_, as the simulation results for *β*_1_ are similar.

## RESULTS

### Application to OHDSI Data

The summaries of patients’ characteristics of the four datasets are listed in **Table 1.** The overall event rates are below 1% for both outcomes. The Medicare data (i.e., MDCR) were observed to have more elderly patients, and therefore a higher prevalence for both the outcome and risk factors of interest.

**Table 1.** Characteristics of the four claims datasets at OHDSI

| Dataset | CCAE | MDCD | MDCR | Optum |
| --- | --- | --- | --- | --- |
| Number of subjects | 64,222 | 59,861 | 69,164 | 62,348 |
| Median Age | 43 | 35 | 71 | 47 |
| % of Female | 69.21 | 73.82 | 68.08 | 69.68 |
| % of Congestive heart failure | 0.70 | 3.06 | 7.58 | 2.61 |
| % of Hypertensive disorder | 20.81 | 31.80 | 57.70 | 32.96 |
| % of Ischemic heart disease | 1.70 | 3.82 | 10.27 | 4.10 |
| % of Type 2 diabetes mellitus | 7.49 | 14.63 | 21.83 | 12.71 |
| % of Coronary arteriosclerosis | 2.39 | 4.92 | 18.43 | 5.75 |
| % of Renal failure syndrome | 0.69 | 2.67 | 2.31 | 2.49 |
| % of Transient cerebral ischemia | 0.41 | 0.64 | 2.32 | 0.71 |
| % of Hyperlipidemia | 20.96 | 22.00 | 43.21 | 33.85 |
| % of Obesity | 7.15 | 16.54 | 6.71 | 9.62 |
| % of Alcohol dependence | 1.79 | 2.94 | 1.01 | 2.29 |
| % of Major depressive disorder | 4.17 | 3.55 | 3.16 | 3.34 |
| % of Acute myocardial infarction | 0.26 | 0.75 | 2.03 | 0.51 |
| % of Stroke | 0.24 | 0.73 | 1.75 | 0.58 |

\*The four claims datasets are CCAE (IBM MarketScan® Commercial), MDCD (IBM MarketScan® Medicaid), MDCR (IBM MarketScan® Medicare) and Optum (Optum© De-Identified Clinformatics).

**Figure 3** shows the estimated log hazard ratios from the three methods as well as their 95% confidence intervals. The ODAC provided hazard ratio estimates are nearly identical to the pooled estimates for most of the risk factors, while meta-analysis estimates have substantial biases compared to the pooled estimator. The differences in estimated effect sizes can lead to potentially different conclusions in the investigation of risk factors. For example, the estimated log hazard ratio of alcohol dependence for AMI changed signs when comparing the pooled estimate (0.571; 95% CI: (0.322, 0.820)) to the meta-analysis estimate (−0.053, 95% CI: (−0.361, 0.255)). In the hazard ratio scale, the pooled estimates was 1.770 (1.380, 2.271) and the meta-analysis estimate was 0.948(0.697, 1.290). Furthermore, the meta-analysis estimates were sometimes qualitatively different from the pooled analysis and ODAC estimates. For example, both the pooled and the ODAC estimates suggested that alcohol dependence is significantly associated with the time to AMI, while the meta-analysis estimate showed it is not a significant risk factor. Similar contrasts were also present regarding risk factors such as hyperlipidemia, obesity and depression. For all the risk factors of stroke, the relative bias of ODAC was less than 2%, whereas the relative bias of meta-analysis estimates were as high as 109%. This real world example suggests that the ODAC estimator is preferred over meta-analysis in the investigation of risk factors of rare time to event outcomes.

**Figure 3:**
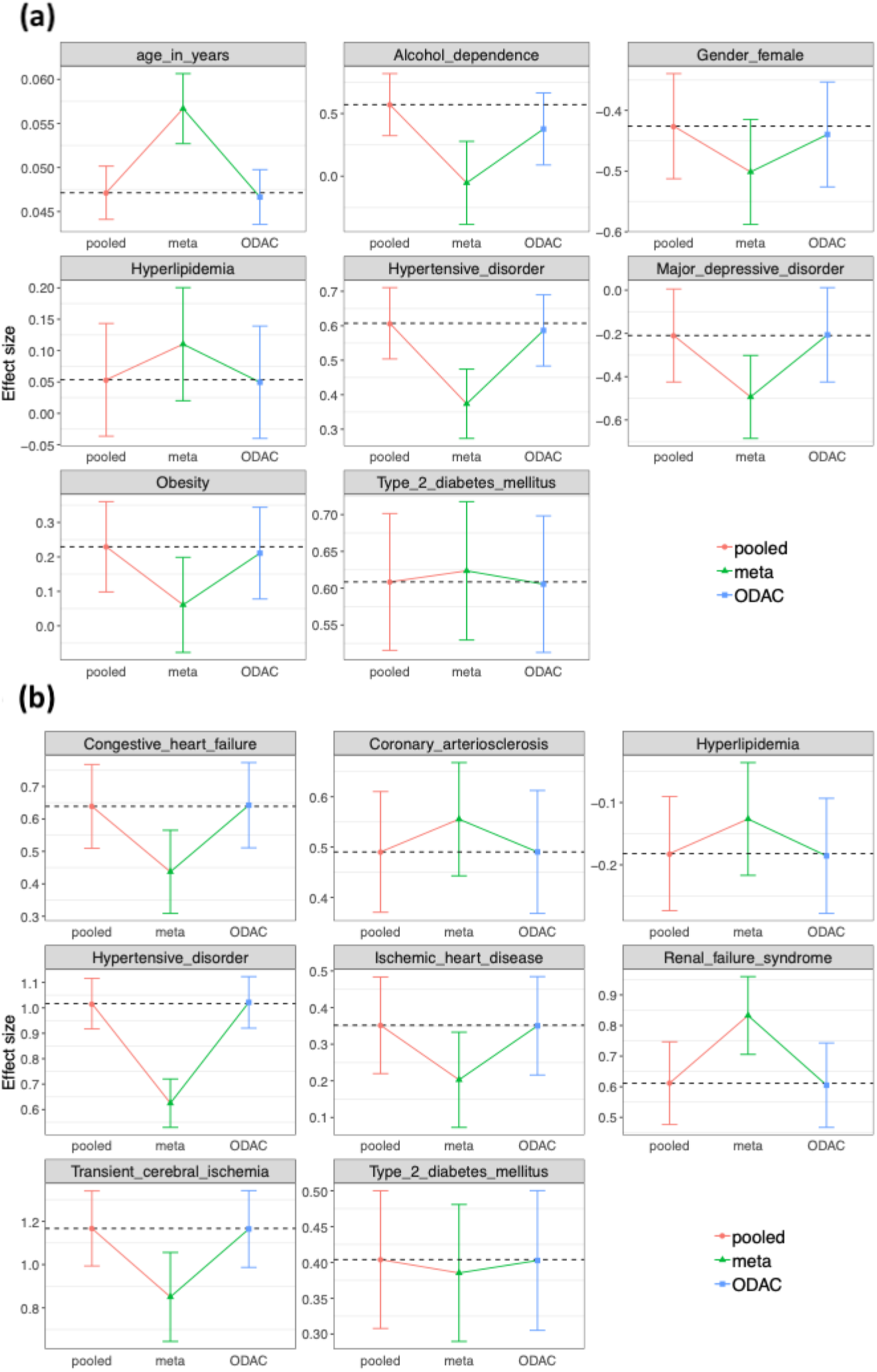
**(a)** Estimated log hazard ratio with 95% confidence intervals for risk factors of AMI using pooled analysis (red), meta-analysis (green), and ODAC (blue); **(b)** Estimated log hazard ratio with 95% confidence intervals for risk factors of stroke using pooled analysis on the combined dataset across all sites (red), meta-analysis (green), and ODAC (blue).

### Numerical expiriments

Our numerical study shows that ODAC has better performance than the meta-analysis estimator especially when the event is rare. The conclusion is supported by **Figure 4,** which shows the box plots of the bias to the pooled estimator for different event rates and sample sizes. We observe that for all scenarios, ODAC obtains relative biases close to 0, meaning that it provides almost identical results to the pooled estimator. When the event rate is fixed and the sample size changes, the performance of the ODAC and meta-analysis estimators stay unchanged. When the event rate decreases, meta-analysis estimator is observed to have larger bias. When event rate is 1%, the average bias of the meta-analysis estimator is around −0.14, while the largest absolute bias of ODAC is 0.01. In addition to the difference in magnitude of bias, the variation of the meta-analysis estimator is much larger compared to that of the ODAC estimator.

**Figure 4:**
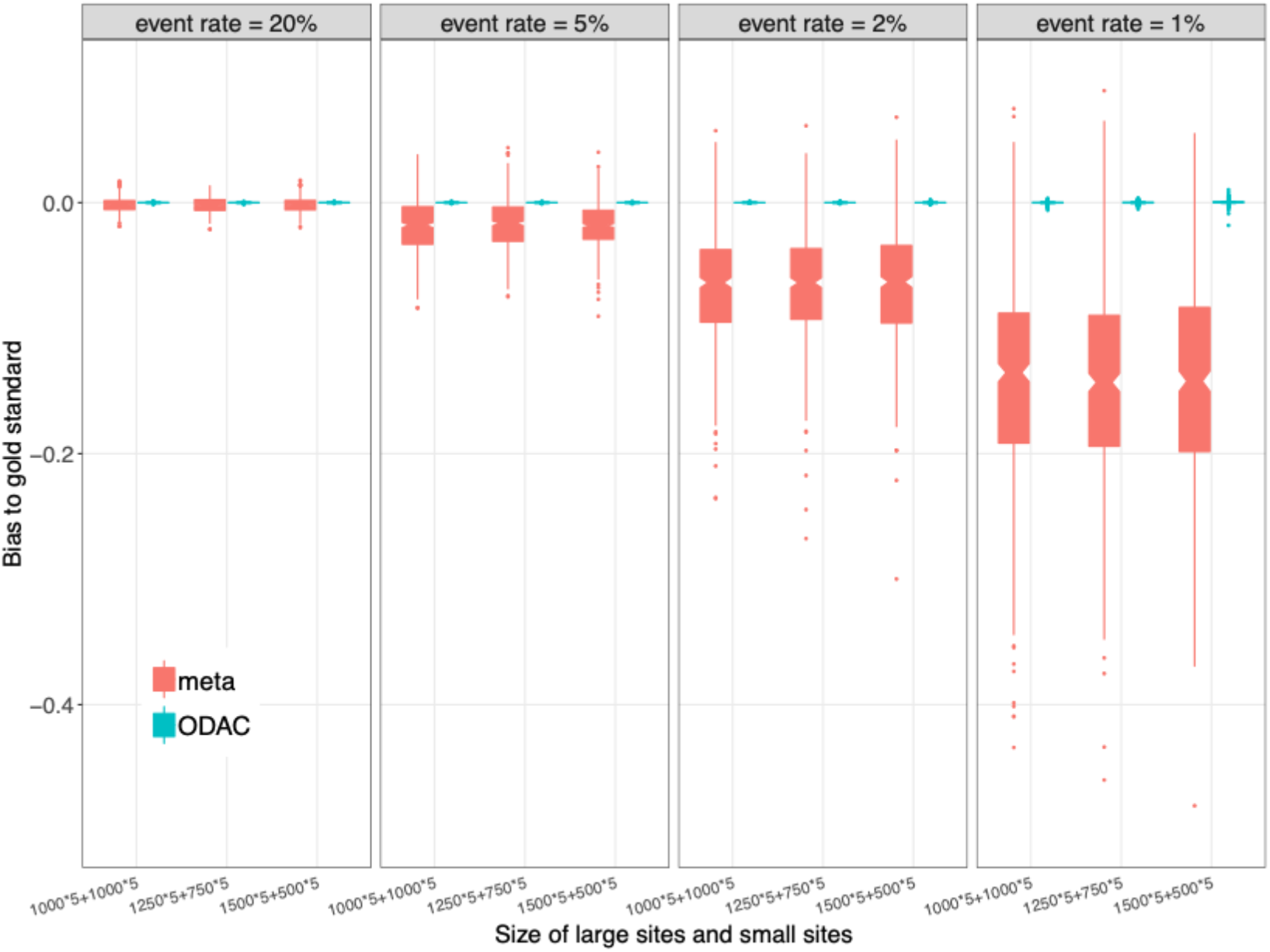
Box plot of relative bias to the gold standard (pooled estimator obtained by fitting Cox model on the combined dataset across all sites) in different simulation settings. The two methods illustrated in the plot are meta-analysis (red) and ODAC (blue). The event rate varies from 20% to 1%, and there are three different sample size distributions: (1) all 10 sites have 1000 samples, (2) five sites with 1250 samples and five with 750 samples; (3) five sites with 1500 samples and five with 500 samples. Under each setting, the boxplots are based on 1000 replications of the experiment.

## DISCUSSION

In this paper, we developed and evaluated ODAC, a distributed, privacy-preserving algorithm to fit a Cox model in a federated multi-center setting. ODAC does not require iterative communication across sites, which minimizes the communication cost, and therefore is easy to implement in large research networks such as OHDSI and PCORnet as well as smaller networks such as those within PCORnet, e.g., PEDSnet (A National Pediatric Learning Health System) [24, 25] and OneFlorida [26].

ODAC was shown to be significantly more accurate than conventional meta-analysis, particular when event rates were low, 1% or below; even though the communication cost (defined as the quantity of numbers needed to be transferred or shared) is greater than that of meta-analysis. Specifically, for a model with *p* parameters, the meta-analysis requires each site to share 2p numbers (a point estimate and a standard error for each parameter), while ODAC requires transferring (*M_t_* + 1)(*p^2^* + *p* + 1) numbers from each site, where *M_t_* is the number of unique event time points across all sites. Usually *M_t_* is not very large in the EHR data setting as the event time is often measured in days. For example, for a one-year follow-up, the largest possible *M_t_* = min (365, *n_e_*), where *n_e_* is the total number of events. When studying extremely rare events, *n_e_* is likely to be much smaller than the number of days during the follow-up period. As a consequence, when the number of covariates in the model is not extremely large, transferring *O*(*M_t_p^2^*) numbers is not considered a burden in multicenter analysis. Yet, the improvement in estimation accuracy is substantial, compared to meta-analysis, particularly when the event rate is low. Compared to iterative methods such as WebDISCO, the communication cost of ODAC is low.

Implementing ODAL in distributive networks such as OHDSI is easy as it does not require iterative communications. The initialization step is fairly standard in a multicenter analysis where each site shares their local estimates. After the initialization, each site is required to share the intermediate quantities *U_k_,Z_k_, W_k_* to all the other sites. These quantities all have closed forms and can be obtained using prewritten code/software packages distributed across the network. Both OHDSI and PCORnet have the necessary infrastructure—ARACHNE (A distributed OHDSI research network and study workflow orchestration) [27] and the PCORnet Query Tool [28] based on PopMedNet (an open-source application used to facilitate multi-site health data networks) [29]to distribute these analytical codes. The surrogate partial likelihood estimator can also be obtained using an optimization function that is commonly found in prewritten code or software packages. The code for ODAC is available from the authors upon request.

The impact of this work is clear, on several dimensions. First, ODAC affords researchers the ability to conduct Cox analyses across many sites in a federated network. Second, ODAC provides a means to preserve the privacy and confidentiality of patients in the context of research conducted on federated data networks. Rather than require individual patient-level data from all network sites, such data are required from only one site. The other sites a proxied by aggregate data from each of the other sites. Third, federated data networks are essential to the study of rare diseases. ODAC estimators were shown to be superior to meta-analysis across a range of sampling regimes including rare outcomes.

Our ODAC algorithm is based on the pooled analysis, which is to fit a unified Cox regression model on the combined dataset. Therefore it essentially requires the data are homogenous distributed across sites. In the future, we plan to extend our method to handle heterogeneity across clinical sites by allowing site-specific effects and covariates. We are also developing open-source software packages to support studies that wish to use ODAC for multicenter analysis in existing distributed networks such as OHDSI and PCORnet. One key task is to implement ODAC according to the networks’ OMOP CDM for OHDSI and the PCORnet CDM to alleviate the potentially complicated preprocessing procedures. We believe that ODAC is a significant contribution to the fast-growing distributed research networks with enormous patient-level RWD who are facing data sharing issues and privacy concerns, and ultimately will help facilitate collaborative efforts to investigate risk factors for time to event outcomes in healthcare systems. Some of the numerous applications where it may be a particularly powerful tool is use is in modeling risk factors for disease progression and treatment effectiveness for uncommon and rare diseases. Because it minimizes communication costs, it will also be an excellent tool for international studies that require time-to-event modeling of outcome.

## CONCLUSION

The proposed ODAC algorithm for multicenter Cox model is privacy preserving and non-iterative. Through a simulation study and a real-world use case using OHDSI data, ODAC was shown to have higher estimation accuracy compared to the meta-analysis, especially for studying rare events.

## CONTRIBUTIONSHIP STATEMENT

RD, CL and YC designed methods and experiments; MJS provided the dataset from OHDSI; CL and JT conducted simulation experiments; MJS conducted data analysis; all authors interpreted the results and provided instructive comments; RD, CL and YC drafted the main manuscript. All authors have approved the manuscript.

