## Supplementary Material for "Learning from local to global - an efficient distributed algorithm for modeling time-to-event data"

This Supplementary Material contains technical details of ODAC.

***Section A: some definitions***

1. The local first order gradient $\nabla L_j(\bar{\beta})$

$$\nabla L_j(\bar{\beta}) = \frac{1}{n_j}\sum_{i=1}^{n_j} \delta_{ij}\{x_{ij} - \frac{\sum_{s\in R_j(T_{ij})} \exp(\bar{\beta}^T x_{sj})x_{sj}}{\sum_{s\in R_j(T_{ij})} \exp(\bar{\beta}^T x_{sj})}\} \qquad \text{(S.1)}$$

2. The local second order gradient $\nabla^2 L_j(\bar{\beta})$

$$\nabla^2 L_j(\bar{\beta}) = \frac{1}{n_j}\sum_{i=1}^{n_j} \delta_{ij}[\frac{\left\{\sum_{s\in R_j(T_{ij})} \exp(\bar{\beta}^T x_{sj})x_{sj}\right\}^{\otimes 2}}{\left\{\sum_{s\in R_j(T_{ij})} \exp(\bar{\beta}^T x_{sj})\right\}^2} - \frac{-\sum_{s\in R_j(T_{ij})} \exp(\bar{\beta}^T x_{sj})x_{sj}^{\otimes 2}}{\sum_{s\in R_j(T_{ij})} \exp(\bar{\beta}^T x_{sj})}] \qquad \text{(S.2)}$$

where $a^{\otimes 2}$ for a vector $a$ denotes the outer product $aa^T$.

3. The global first order gradient $\nabla L(\bar{\beta})$

$$\nabla L(\bar{\beta}) = \frac{1}{N}\sum_{j=1}^{K}\sum_{i=1}^{n_j} \delta_{ij}\{x_{ij} - \frac{\sum_{(s,k)\in R_{ij}} \exp(\bar{\beta}^T x_{sk})x_{sk}}{\sum_{(s,k)\in R_{ij}} \exp(\bar{\beta}^T x_{sk})}\} \qquad \text{(S.3)}$$

4. The global second order gradient $\nabla L(\bar{\beta})$

$$\nabla^2 L(\bar{\beta}) = \frac{1}{N}\sum_{j=1}^{K}\sum_{i=1}^{n_j} \delta_{ij}\frac{\left\{\sum_{(s,k)\in R_{ij}} \exp(\bar{\beta}^T x_{sk})x_{sk}\right\}^{\otimes 2} - \sum_{(s,k)\in R_{ij}} \exp(\bar{\beta}^T x_{sk})x_{sk}^{\otimes 2}}{\left\{\sum_{(s,k)\in R_{ij}} \exp(\bar{\beta}^T x_{sj})\right\}^2} \qquad \text{(S.4)}$$

*Section B: Distributive calculation of $\nabla L(\bar{\beta})$ and $\nabla^2 L(\bar{\beta})$*

**Step 1**: Calculate the terms $\sum_{(s,k)\in R_{ij}} exp(\bar{\beta}^T x_{sk})$, $\sum_{(s,k)\in R_{ij}} exp(\bar{\beta}^T x_{sk}) x_{sk}$, and $\sum_{(s,k)\in R_{ij}} exp(\bar{\beta}^T x_{sk}) x_{sk}^{\otimes 2}$ distributively using the aggregated information $\{U_k, W_k, Z_k\}$:

Since there is a factor $\delta_{ij}$ in the equation, we only need to consider the time point $T_{ij}$ where $\delta_{ij} = 1$, i.e, the combined event time set, $\mathcal{T} = \cup_{ij} \{T_{ij}: \delta_{ij} = 1\}$. We can also sort $\mathcal{T} = \{t_1 \le t_2 \ldots \le t_d\}$,

and for any $T_{ij}$ such that $\delta_{ij} = 1$, there is some $r \in \{1, \ldots, d\}$, such that $T_{ij} = t_r$. Then we have $R_{ij} = R(t_r)$, and

$$\sum\nolimits_{(s,k)\in R_{ij}} exp(\bar{\beta}^T x_{sk}) = \sum\nolimits_{(s,k)\in R(t_r)} exp(\bar{\beta}^T x_{sk}) x_{sk} = \sum_{k=1}^{K} U_k(t_r),$$

$$\sum\nolimits_{(s,k)\in R_{ij}} exp(\bar{\beta}^T x_{sk}) x_{sk} = \sum\nolimits_{(s,k)\in R(t_r)} exp(\bar{\beta}^T x_{sk}) x_{sk} = \sum_{k=1}^{K} W_k(t_r)$$

and

$$\sum\nolimits_{(s,k)\in R_{ij}} exp(\bar{\beta}^T x_{sk}) x_{sk}^{\otimes 2} = \sum\nolimits_{(s,k)\in R(t_r)} exp(\bar{\beta}^T x_{sk}) x_{sk}^{\otimes 2} = \sum_{k=1}^{K} Z_k(t_r)$$

where the terms $U_k = (U_k(t_1), U_k(t_2), \ldots, U_k(t_d))$, $W_k = (W_k(t_1), W_k(t_2), \ldots, W_k(t_d))$ and $Z_k = (Z_k(t_1), Z_k(t_2), \ldots, Z_k(t_d))$ have been broadcasted such that it is accessible to all site.

**Step 2**: Distributively calculate $\nabla L(\bar{\beta})$ and $\nabla^2 L(\bar{\beta})$:

After calculating the terms $\sum_{(s,k)\in R_{ij}} exp(\bar{\beta}^T x_{sk})$, $\sum_{(s,k)\in R_{ij}} exp(\bar{\beta}^T x_{sk}) x_{sk}$, and $\sum_{(s,k)\in R_{ij}} exp(\bar{\beta}^T x_{sk}) x_{sk}^{\otimes 2}$ in each site, we can distributively calculate $\nabla L(\bar{\beta})$ and $\nabla^2 L(\bar{\beta})$ by

$$\nabla L(\bar{\beta}) = \frac{1}{N}\sum_{j=1}^{K} f_k(\bar{\beta}),$$

where

$$f_k(\bar{\beta}) = \sum_{i=1}^{n_j} \delta_{ij}\{x_{ij} - \frac{\sum_{(s,k)\in R_{ij}} exp(\bar{\beta}^T x_{sk}) x_{sk}}{\sum_{(s,k)\in R_{ij}} exp(\bar{\beta}^T x_{sk})}\},$$

and

$$\nabla^2 L(\bar{\beta}) = \frac{1}{N}\sum_{j=1}^{K} g_k(\bar{\beta}),$$

where

$$g_k(\bar{\beta}) = \sum_{i=1}^{n_j} \delta_{ij} \frac{\left\{\sum_{(s,k)\in R_{ij}} exp(\bar{\beta}^T x_{sk})\, x_{sk}\right\}^{\otimes 2} - \sum_{(s,k)\in R_{ij}} exp(\bar{\beta}^T x_{sk})\, x_{sk}^{\otimes 2}}{\left\{\sum_{(s,k)\in R_{ij}} exp(\bar{\beta}^T x_{sj})\right\}^2}.$$

The term $\mathbf{f_k}(\bar{\boldsymbol{\beta}})$ and $\mathbf{g_k}(\bar{\boldsymbol{\beta}})$ can be calculated locally without patient-level data from other sites.

#### *Section C: Variance estimator of ODAC*

The variance of $\tilde{\beta}_j$, denoted by $\tilde{V}_j$ can be estimated by

$$\tilde{V}_j = \frac{1}{N}\left\{-\nabla^2 L_j(\tilde{\beta}_j)\right\}^{-1}, \qquad \text{(S.5)}$$

by plugging in $\tilde{\beta}_j$ in (S.2).

The variance of $\tilde{\beta}$ denoted by $\tilde{V}$ can be estimated by

$$\tilde{V}_j = \frac{1}{N}\left\{-\nabla^2 L(\tilde{\beta})\right\}^{-1}, \qquad \text{(S.6)}$$

by plugging in $\tilde{\beta}$ in (S.4).

### *Section D: Additional numerical studies*

#### *Bias to the true parameter value*

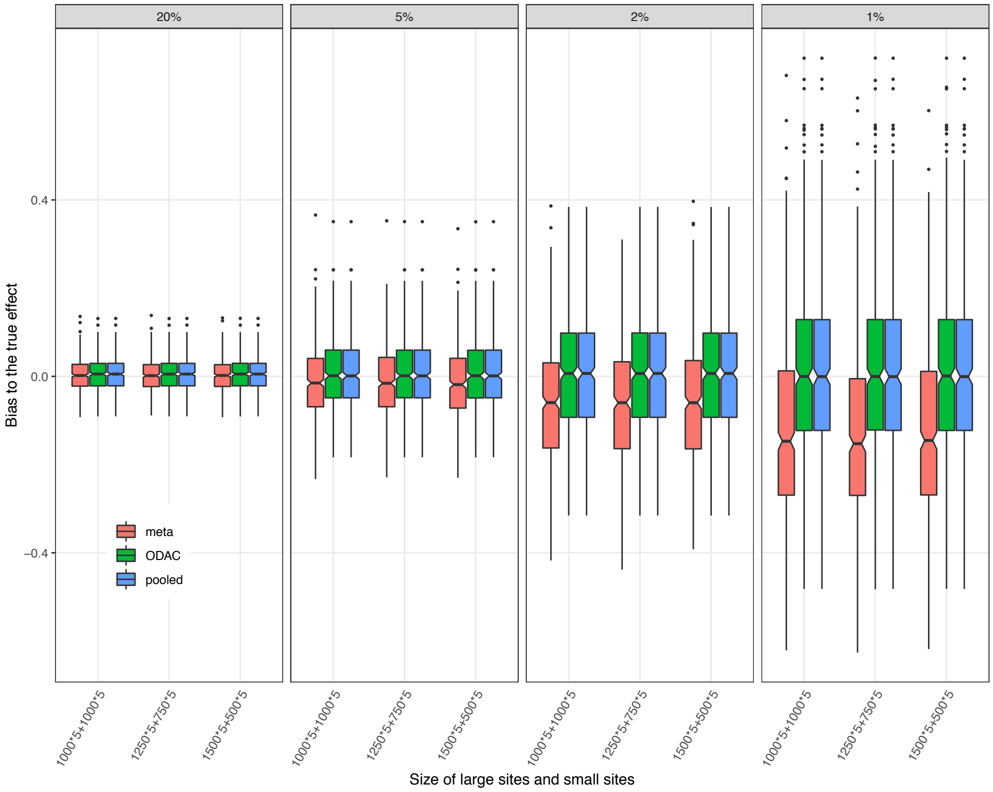


Figure S1. Box plot of bias to the true parameter value in different simulation settings. The three methods illustrated in the plot are meta-analysis (red) and ODAC (green), and the pooled analysis (blue). The event rate varies from 20% to 1%, and there are three different sample size distributions: (1) all 10 sites have 1000 samples, (2) five sites with 1250 samples and five with 750 samples; (3) five sites with 1500 samples and five with 500 samples. Under each setting, the boxplots are based on 1000 replications of the experiment.

We considered the same setting as the main paper, where a pooled dataset of N=10,000 subjects were generated based on Weibull proportional hazard model, where the baseline hazard follows a

Weibull-distribution with scale 200 and shape 20. We generated two covariates from i.i.d. uniform distributions and the true log hazard ratios were set to be $\beta$= (-1, 2). We set the event rate (number of cases over number of subjects) as 20%, 5%, 2% and 1% by appropriately modifying the distribution of censoring times. The pooled data were distributed over K=10 clinical sites, with five large and five small sites. We set the relative sizes of the large to small sites to be 1000/1000, 1250/750, 1500/500. Under each scenario, we compared the ODAC estimator to the meta-analysis estimator over 1000 replications. From Figure S1, From the figure, we can see that the bias for meta-analysis is increasing when the event rate is decreasing. However, ODAC is able to provide similar results as the pooled analysis.

***Evaluation by subsetting a real dataset***

We use the lung cancer dataset from the North Central Cancer Treatment Group (NCCTG), which contains 228 observations with variables such as survival time, censoring indicator and other covariates such as age, sex, etc. For more information about the dataset, please see the link (https://stat.ethz.ch/R-manual/R-devel/library/survival/html/lung.html). We split the dataset randomly into four smaller datasets. We ran ODAC, meta-analysis and the pooled analysis on these four smaller datasets using a Cox model with age and sex as covariates. We repeat the above procedure 100 times and use the following boxplot to show the estimates over the 100 replications. From the Figure S2, we see that ODAC is much more stable compared to meta-analysis and provides almost identical results as the pooled analysis, which further demonstrated the benefit of ODAC compared to meta-analysis.

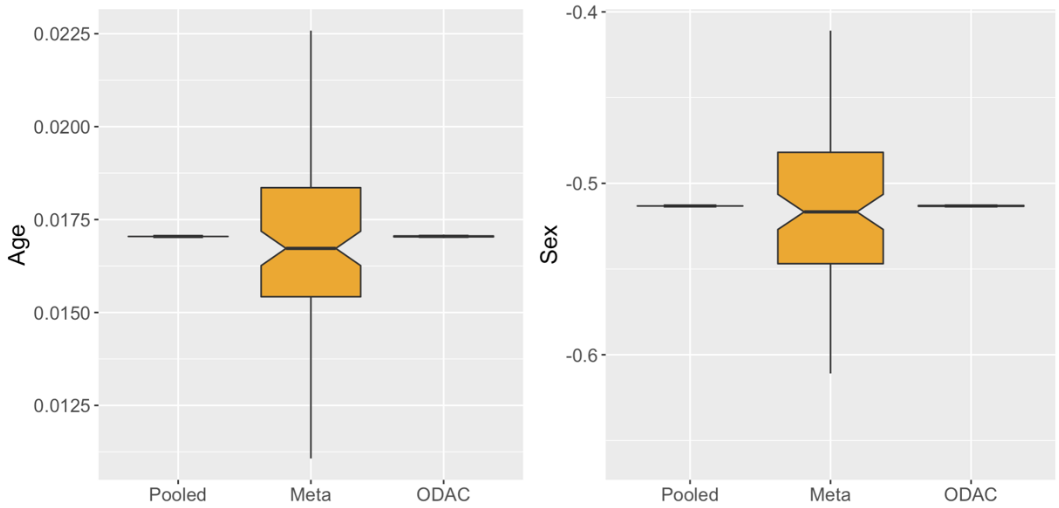


Figure S2: Boxplot of estimates from the pooled analysis, meta-analysis and ODAC using a lung cancer dataset. The dataset is split randomly into four subsets. The boxplot is based on 100 replications. The pooled analysis has identical results across the 100 replications since it is obtained from the whole dataset. ODAC provides similar results as the pooled analysis, while meta-analysis has larger variation.
